# Type II photosynthetic reaction center genes of avocado (*Persea americana* Mill.) bark microbial communities are dominated by aerobic anoxygenic Alphaproteobacteria

**DOI:** 10.1101/2020.09.12.295014

**Authors:** Eneas Aguirre-von-Wobeser

## Abstract

The tree bark environment is an important microbial habitat distributed worldwide on thrillions of trees. However, the microbial communities of tree bark are largely unknown, with most studies on plant aerial surfaces focused on the leaves. Recently, we presented a metagenomic study of bark microbial communities from avocado. In these communities, oxygenic and anoxygenic photosynthesis genes were very abundant, especially when compared to rhizospheric soil from the same trees. In this work, Evolutionary Placement Algorithm analysis was performed on metagenomic reads orthologous to the *PufLM* gene cluster, encoding for the bacterial type II photosynthetic reaction center. These photosynthetic genes were found affiliated to different groups of bacteria, mostly aerobic anoxygenic photosynthetic Alphaproteobacteria, including *Sphingomonas, Methylobacterium* and several Rhodospirillales. These results suggest that anoxygenic photosynthesis in avocado bark microbial communities functions primarily as additional energy source for heterotrophic growth. Together with our previous results, showing a large abundance of cyanobacteria in these communities, a picture emerges of the tree holobiont, where light penetrating the trees canopies and reaching the inner stems, including the trunk, is probably utilized by cyanobacteria for oxygenic photosynthesis, and the far-red light aids the growth of aerobic anoxygenic photosynthetic bacteria.

## Introduction

There are trillions of trees growing worldwide [1], constituting an important habitat for epiphytic bacteria. However, most of the attention to tree aerial habitat has been centered on the leaves, with stem bark as a microbial habitat largely neglected [2]. Recently, 16S microbiome surveys of bark from different woody plants have revealed bark surfaces as important microbial environments, usually more diverse than the leaves [3-7].

Recently, a bark metagenomic study was conducted on bark from avocado trees, revealing highly diverse microbial communities dominated by bacteria, with the presence of archaea, fungi, algae and other eukaryotic groups [8]. Functional analysis of these communities revealed the presence of many oxygenic and anoxygenic photosynthetic genes, which were absent from rhizospheric soil from the same trees [9]. The presence of these genes in avocado bark suggests active phototrophic communities. However, a detailed phylogenetic analysis is needed to better understand the bacterial groups potentially capable of anoxygenic photosynthesis in these communities.

Phylogenetic analysis of metagenomic sequences is challenging, as many non-overlapping fragments are found of a potentially diverse collection of othologs for a given gene. This complicates the correct alignment of reads, and can result in biases during tree reconstruction. Atamna-Ismaeel et al. [10] reconstructed phylogenetic trees from the genes encoding for the type II photosynthetic reaction center, PufL and PufM for a collection of different metagenomic data-sets, including several from plant leaves. By using several datasets, and focusing in a fragment from each gene, they managed to get enough coverage for alignment and three inference. However, this approach proved difficult to implement for a single metagenomic dataset.

Recently, Evolutionary Placement Algorithms (EPA) have been developed, in which a reference tree is constructed from known sequences of the gene of interest, and the most likely position of each query metagenomic read is found in the branches of that tree [11]. These algorithms have many advantages, as any number of reads can be mapped (or placed, as it is formally termed) to the tree. Furthermore, reads are not only placed at the branches of the tree, but they can find their position at the inner branches. These inner branch placements can be interpreted as phylogenetic relations with the taxa at the tips, equivalent to new bifurcating branches in a traditional phylogenetic tree. The deeper a placement is observed, the more ancestral its relation with the taxa at the tips.

In this work, the diversity and phylogenetic affiliation of anoxygenic photosynthetic bacteria utilizing type II reaction centers in avocado bark microbial communities is explored. EPA analysis was performed on the avocado bark metagenomic dataset presented previously [8, 9] of the *PufLM* gene cluster, encoding for a type II photosynthetic reaction center. A dominance of aerobic anoxygenic photosynthetic bacteria was clearly observed among *PufLM-*carrying bacteria of the avocado bark environment.

## Methods

### Metagenomic dataset

The analysis presented in this work is built on a database of metagenomic data from Hass variety avocado tree bark, which was presented before [8]. Here, I present an analysis performed on DNA sequences from an organic-managed orchard located near Malinalco, Mexico. Details on the orchard can be found at Aguirre-von-Wobeser *et al*., [8]. Approximately 5 x 5 cm incisions were collected at about 1 m from the ground on the main stem (trunk) of avocado trees. The periderm was carefully removed and subjected to DNA extraction using a PowerSoil extraction kit (Qiagen, USA). Three bark samples from the organic orchard yielded high-quality DNA, and where used for this analysis. Extracted DNA was used for library preparation with a DNA Flex kit (Illumina, USA) and sequenced on a nextSeq machine (Illumina).

### Identification of PufLM reads

*PufLM* fragments were identified in the metagenomic reads from avocado bark using the blastx algorithm, run with Diamond [12], using a e-value threshold of 0.001. Two collections of *PufLM* sequences were used as references for this step. First, a comprehensive *PufLM* database from Imhoff et al. [13] was used, which contains well-characterized *PufLM* genes from genome-sequencing projects, most of which were derived from type strains. This database contains 197 full-length sequences of concatenated *PufL* and *PufM* genes, and is representative of the current knowledge on the diversity of these genes, covering strains with a wide range of phylogenetic affiliations and environmental origins. Second, a database which was originally derived from environmental samples through metagenomics from phyllospheres of trees was used [10]. This database has fractions of the pufL and pufM genes in separate files. Therefore, the reads identified with blastx using these references correspond to matches restricted to these fractions.

### Phylogenetic placement

All reads identified as fractions of the *PufL* and *PufM* reads were placed on a phylogenetic tree of the Imhoff et al. [13] database, for which they were previously put in the adequate reading frame (reverse complement as necessary) and translated to amino acids. To build the reference tree, the *PufLM* sequences were re-aligned using Muscle [14], resulting in an alignment almost identical as in the original database. This alignment was used for phylogenetic tree reconstruction with RAxML [15]. The most adequate substitution model for these sequences was determined by RAxML to be LG.

The metagenomic reads were aligned to the reference alignment using PaPaRa [16], resulting in a single file with the reference and query sequences aligned. This alignment, and the reference phylogenetic tree, were used as inputs for the placement algorithm, which was run with the option -f v of RAxML [17]. The resulting placement was explored and visualized with functions from the Gappa package [18].

### Data availability

All sequences analyzed in this study have been previously presented and are available at NCBI Short Read Archive (BioProject PRJNA656796, accession SUB7881803) and KBase (https://kbase.us/n/69195/32/).

## Results

### Localization of PufLM genes in Bark metagenomic data

Metagenomic reads corresponding to bacterial reaction center type II (*pufLM* genes) were located in metagenomic reads from avocado bark samples using blast methods. Two data-sets were used as references for this purpose. The Imhoff et al. [13] reference contained genes from carefully selected strains, including type strains when available, representatives of other strains from groups lacking type materials, and some strains of interest for the study of prokaryotic photosynthesis. The Atamna-Ismaeel et al. [10] reference contained *PumL* and *PufM* genes recovered from strains from the leaves of several, mostly herbaceus plants. We recovered a total of 2400 *pufLM* metagenomic reads from avocado bark using the Imhoff et al. [13] reference and 4082 using the Atamna-Ismaeel et al. [10] reference (Table 1).

**Table 1.** PufML reads recovered from the avocado bark metagenomic dataset

| Sample | Imhoff et al. (2018)<br>reference (PufLM) | Atamna-Ismaeel et al.<br>(2012) reference (PufL) | Atamna-Ismaeel et al.<br>(2012) reference (PufM) |
| --- | --- | --- | --- |
| A1 | 580 | 439 | 497 |
| A4 | 862 | 701 | 789 |
| A5 | 958 | 767 | 889 |

### Phylogenetic placement of PufLM reads

Evolutionary phylogenetic placement was used to map the short metagenomic reads from avocado bark to the branches of a reference phylogenetic tree. In order to obtain well-characterized phylogenetic affiliations of the avocado bark anoxygenic photosynthetic bacteria, the Imhoff et al. [13] phylogenetic tree was used as a reference for placement. First, the Imhoff et al. [13] sequences were re-aligned, and the phylogenetic tree was reconstructed using RAxML. The obtained tree had essentially the same topology as the published tree from these sequences [13], indicating a robust reconstruction of the phylogenetic relationships of these genes (Figure 1). Evolutionary phylogenetic placement indicated the affiliation of the avocado bark reads to known *PufLM* reads. The obtained placements were essentially the same with reads collected using the Imhoff et al. [13] reference (Figure 1) and the *Atamna-Ismaeel reference (Figure S1). Most avocado bark PufLM genes were affiliated to a restricted set of branches from the known PufLM phylogeny, including the Sphingomonadales, different groups fo Rhizobiales, and specific branches of Rhodospirillales and Chloroflexales*.

**Fig. 1.**
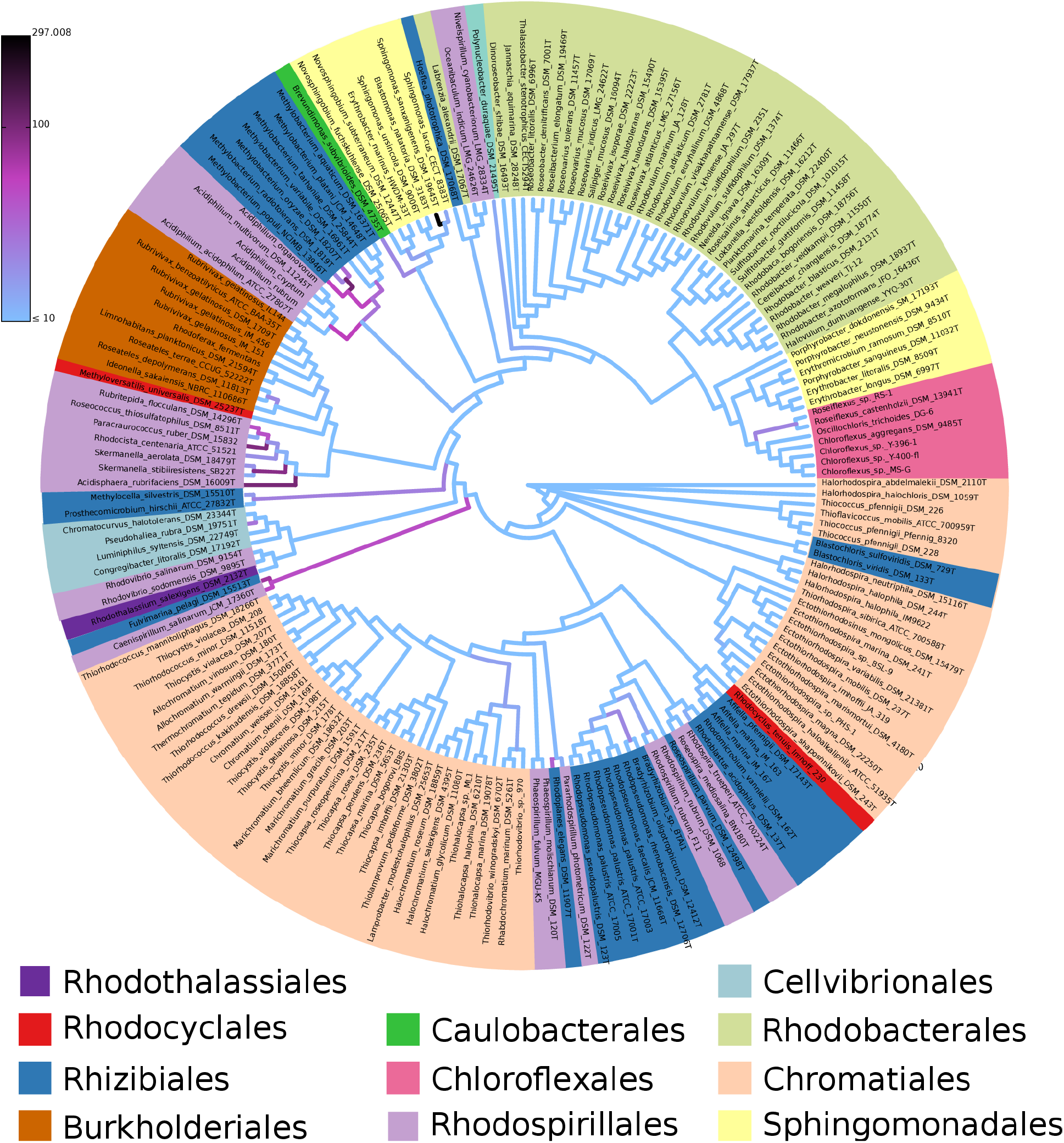
Evolutionary Placement of metagenomic reads from avocado (*Persea americana*) bark on a PufLM tree reconstructed from the sequences presented in Imhoff et al. (2018). The metagenomic reads where selected with blast methods using the *PufL* and *PufM* sequences presented in Atamna-Ismaeel et al. (2012). Placements are shown as read counts in the branches of the trees, according to the scale. For the placement, reads from three avocado bark samples were pooled.

### Avocado bark Sphingomonadales

*PufLM* genes belonging to the Sphingomonadales were very abundant in bark microbial communities (Figure 1). All of these genes were affiliated to the Sphingomonadaceae family, with no placements mapping to the Erythrobacteraceae. Particularly, most of the genes fragments from this groups mapped to the tip of the branch corresponding to *Sphingomonas sanxanigenens*, and to a much lower extent to *S. lacus*. Although the exact species assignation of these reads is not certain, as many species are not represented in the reference, placement analysis indicates that close relatives of *S. sanxanigenens* dominate the photosynthetic Sphingomonadales on avocado bark.

Other *PufLM* gene fragments from the Sphingomonadaceae family mapped to the base of the *Novosphingobium* genus branch, including *Erythrobacter marinus*, which is probably misclassified [13]. This suggests the presence of photosynthetic bacteria related to *Novosphingobium*, belonging to species not included in the reference sequences.

### Avocado bark Rhizobiales

Many reads from avocado bark microbial communities where mapped to different Rhizobiales groups by the placement algorithm. Of these, one branch of *Methylobacterium*, which included *M. oryzae, M. populi* and *Methylobacterium radiotolerans* had many placements at the tips, as well in internal branches. This shows that avocado bark communities include a diversity of species related to this group. Two of these species, *M. oryzae* and *M. populi* where originally isolated from plants [19-20], suggesting that they are common inhabitants of plant surfaces. Curiously, *PufLM* genes from another *Methylobacterium* branch, including *M. aquaticum, M. platani, M. tarhaniae* and *M. variabile* were absent from avocado bark in this study. From these, *M. platani* was isolated from a leaf of a *Platanus orientalis* tree [21](Kang et al., 2007), and the other three from water or soil. Only the basal branch leading to this *Methylobacterium* cluster showed placements, indicating the presence of a related, yet independent group in avocado bark.

Other Rhrizobiales groups had *PufLM* genes in the bark envrionment. These genes do not form a cohesive phylogenetic group [13], and are found in different branches of the *PufLM* grene phylogeny (Figure 1). The *Blastochloris* genus had placements in a long branch starting deep in the phylogenetic tree, but not in the tips. The separation of *Blastochloris* from other Rhizobacteria has been explained [13], since this genus uses bacteriochlorophyll *b* instead of bacteriocholophyll *a* in its reaction center [22]. The placement of metagenomic reads at the basal branch of this group suggests that other lineages, related to the *Blastochloris* references, but with different *PufLM* genes, exist in the bark communities of avocado. Methagenomic reads were also mapped to *Methylocella sylvestris*, both at the tip and at a branch connecting this species with Prosthecomicrobium hirschii. *Methylocella sylvestris* is a methanotrophic bacterium [23]. *Fulvimarina pelagi*, antoher rhizobacterium, and *Caenispirillum salinarum* of the order Rhodospirillales, which PufLM genes clustered tightly [13], were also represented by placements at both tips and at the branch connecting them. Placements were also observed at two inner branches of the *Bradyrhizobium/Rhodopseudomonas* cluster, suggesting the presence of *PufLM* genes from different species of those genera, or genera related to them.

Genes for the photosynthtetic type II reaction center were also found for the order Rhodospirillales. The group containing *Roseococcus, Rubritepida, Skermanella*. *Paracraurococcus, Rhodocista* and *Acidisphaera* had many *PufLM* placements, both at the tips and at internal branches.

Another group that showed many *PufLM* genes in the avocado bark samples included an internal branch joining two species of the genus *Roseiflexus* (Chloroflexales). In the reads recovered with the Atamna-Ismaeel references, his group also had many more placements in internal branches and at the tips (Figure S1).

## Discussion

Metagenomic read placement of avocado bark *PufLM* reads was highly concentrated on the *Sphingomonas sanxanigenens* tip of the phylogenetic tree. *Sphingomonas* is a highly diverse genus, which includes photosynthetic and non-photosynthetic bacteria. *Sphingomonas* was one of the most abundant and diverse genera in avocado bark microbial communities overall [8]. Moreover, this genus has been found to dominate on bark surfaces of other trees [4, 7], as well as on leafs [24]. The high abundance of *PufLM* genes detected for *Sphingomonas*, particularly mapping to *S. sanxanigenens*, indicates that this species, or closely related species, are major contributors to the anoxygenic photosynthetic potential in this environment. Photosynthetic *Sphingomonas* are among the bacteria capable of performing anoxygenic photosynthesis in the presence of oxygen [13], and they typically require organic carbon substrates for growth. The capability of performing photosynthesis in the presence of oxygen, as an additional source of energy, could be an advantage on plant surfaces including leafs and bark, as they are not restricted to finding and/or creating anoxic niches for photosynthesis. Interestingly, the type strain of *S. sanxanigenens* was isolated from soil from a wheat plantation [25]. Given the high plant biomass found on wheat plantations, and the dominance of *S. saxanigenes* in the bark environment, an association of this species with plants seems likely. Remarkably, the absence fo *PufLM* genes in soil in this study suggests that the main habitat of these bacteria is the surfaces of plants, rather than soil.

Bacteria from the *Methylobacterium* genus have been observed in 16S microbiome studies of bark surfaces from different trees [7], and are very common on leaf surfaces [24]. In a previous analysis, it was shown that *Methylobacterium* is one of most abundant genera in avocado bark microbial communities [8]. *PufLM* genes from *Methylobacterium* were observed mainly in one group of species from the references available for this genus. Within this group, however, reads were placed at branches at many different levels, suggesting a high diversity of *PufLM* genes, as compared to other abundant taxa, like *Sphingomonas*. This suggests a more flexible photosynthtetic apparatus, possibly with more structural diversity in this genus. A characteristic feature of members of *Methylobacterium* is their ability to utilize methanol and other one-carbon molecules for growth. Furthermore, some *Methylobacterum* species can degrade complex organic molecules, including refractory compounds [26, 27]. In the bark environment, they could access difficult to degrade plant-derived molecules as carbon sources. Complete sets of photosynthetic genes are commonly found in *Methylobacterium* [28], and bacteriochlorophyll production in strains isolated from leaves has been demonstrated by far-red light absorption *in vivo* [29]. Therefore, photosynthetic activity in this group is almost certain, however, its physiological role is still being elucidated. A metabolic model of *Methylobacterium extorquens* AM1 found all the genes necessary for carbon fixation through the Calvin Cycle, suggesting the possibility of autotrophic growth [30]. However, it is also possible that CO_2_ fixation is utilized as a mechanism to dispose excess electrons from reduced cofactors [30], as has been shown for other bacteria [31].

The presence of *Blastochloris* reads in bark microbial communities, different from the reference sequences, is interesting, as only four species from this genus are known [22]. Mapping of reads from the avocado bark metagenome to *Methylocella* species *PufLM* suggests the posibility of methanotrophy in this environment. However, no methane monoxygenase genes were fund in any bark sample [9], which casts doubt on the presence of methanotrophic activity in avocado bark.

A group of Rhodospirillales was well represented in the *PufLM* sequences of the avocado bark metagenome. It included many placements in the whole branch, including the tips and internal branches. From these, reads from *Paracraurococcus ruber* and *Acidisphaera rubrifaciens* were especially abundant. There is not much information about these bacterial gropus, and the presence, diversity and significance of these genes in the bark environment should be explored in future studies. The only group of bacteria with abundant *PufLM* genes in avocado bark outside of the class Alphaproteobacteria, was a branch Chloroflexales containing the genus *Roseiflexus*. These species are known from extreme environments [32], and the presence of related genes in the bark environment is interesting. *Roseiflexus* perform photosynthesis under anaerobic light conditions, and can also thrive chemoheterotrophically when oxygen is present. To test whether close relatives of *Roseiflexus* were found in the bark environment, a search for metagenomic reads was conducted in our dataset using the genome of *Roseiflexus* RS-1 as a reference. Only a few matches were found (data not shown), indicating that the homology is restricted to the *PufLM* genes.

Most of the bacterial type II reaction centers found in this study belonged to Aerobic Anoxygenic Photosynthetic Alphaproteobacteria, with a dominance from *Sphingomonas, Methylobacterium*, and a diversity of Rhodospirillales. This suggests that the role of anoxygenic photosynthesis in the bark environment is mainly as an additional energy source for heterotrophic growth. In a previous analysis of the avocado bark metagenome dataset, many genes for cyanobacterial photosynthesis were found [9]. The present study completes the picture of photosynthetic activity in the tree holobiont. Light not harvested in the canopy, and reaching the inner stems of the trees, could be enriched in wavelengths left unused by the leaves. This light seems to support phtotrophic communities, where Cyanobacteria utilize remaining light in the visible spectrum, absorbed by phycobilisomes for carbon fixation [9], and Aerobic Anoxygenic Photosynthetic bacteria utilize far-red light as an additional energy source for photoheterotropic growth.

## Conclusions

Genes for the bacterial anoxygenic type-II photosynthetic apparatus found in avocado bark were mainly from Aerobic Anoxygenic Photosynthetic bacteria. From these, *Sphingomonas* PufLM genes had low diversity as compared to *Methylobacterium*, which where more abundant. PufLM genes from other bacterial groups were present in the avocado bark environment, including Rhodospirillales, *Blastochloris*, and bacteria from the phylum Chloroflexia. In many cases, metagenomic reads mapped to internal branches, indicating the presence of PufLM genes from phyla not represented in the reference sequences. This suggests that unknown groups of photosynthetic bacteria might be present in this microbial environment. The present study confirms that the avocado bark environment is an important habitat for photosynthetic bacteria, which probably thrive on light unutilized by the canopy leaves.

## Acknowledgments

I thank the following people and institutions: Ulricke Wiegel and Jesús Torres for allowing sampling of their avocado trees, and the Mexican Council of Science and Technology (CONACyT) and the Mexican Ministry of Education (SEP) for providing financial support, through grant CB-2014-01-242956.

## Declarations

Conflicts of interest/Competing interests – Non declared.

