## Supplementary materials for "Type II photosynthetic reaction center genes of avocado (*Persea americana* Mill.) bark microbial communities are dominated by aerobic anoxygenic Alphaproteobacteria"

Eneas Aguirre-von-Wobeser

### Supplementary Materials

CONACYT – Centro de Investigación y Desarrollo en Agrobiotecnología Alimentaria, Centro de Investigación y Desarrollo, A.C., Blvd. Sta. Catarina s/n, Col. Santiago Tlapacoya, 42110, San Agustín Tlaxiaca, Hidalgo, Mexico

Correspondance:

E. Aguirre-von-Wobeser

52-55-25-24-86-26

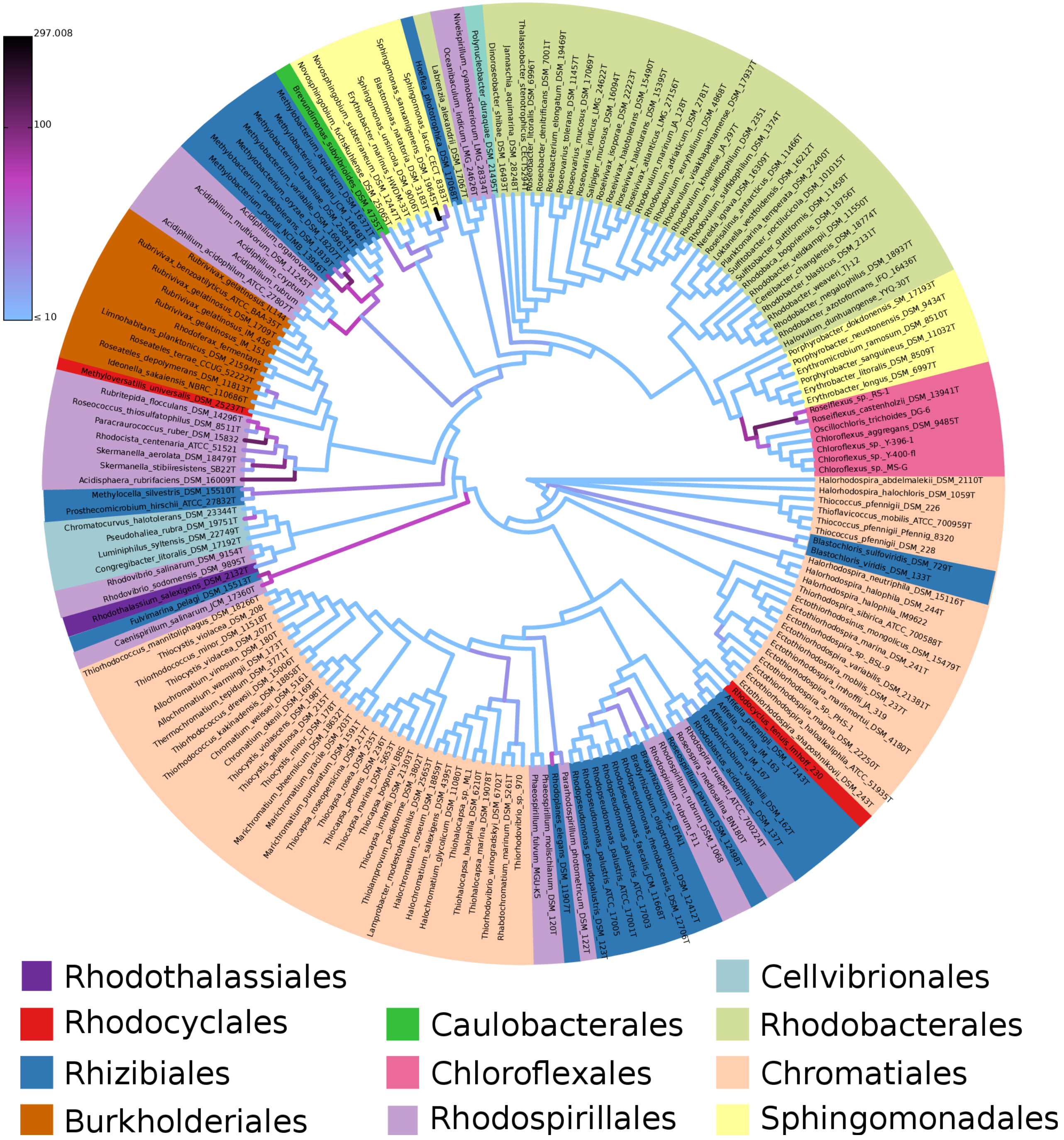


**Fig. S1** Evolutionary Placement of metagenomic reads from avocado (*Persea americana*) bark on a PufLM tree reconstructed from the sequences presented in Imhoff et al. (2018). The metagenomic reads where selected with blast methods using the *PufL* and *PufM* sequences presented in Atamna-Ismaeel et al. (2012). For the placement, reads from three avocado bark samples were pooled.
